# Anellovirus protein coded by ORF2/3 recruits host cell replication and homologous recombination machinery during replication

**DOI:** 10.1101/2025.05.28.656439

**Authors:** Nicole Boisvert, Stephanie Thurmond, Carmen Elenberger, Patricio Jeraldo, Cato Prince, Nolan Sutherland, Jose Melo, Ken Tsheowang, Cameron Dodier, Maciej Nogalski, Dinesh Verma, Geoffrey Parsons, Joseph Cabral

## Abstract

*Anelloviridae* is a family of single-stranded DNA viruses that are thought to be non-pathogenic and commensal. Despite their ubiquitous presence in human populations, little is known about the anellovirus mechanism of replication in host cells. We identified the protein coded by ORF2/3 as necessary and sufficient to initiate replication from the minimal origin of replication for viruses of both the *Beta-* and *Alphatorquevirus* genera. Supporting this observation, we identified components of the polymerase alpha and BTR complexes as interacting with the viral replication initiation protein (Rip) during DNA replication, suggesting a recombination-dependent mechanism of replication that uses host cell machinery to mediate dissolution of replication intermediates. Furthermore, we mapped a 92-bp minimal origin of replication sequence for the *Betatorquevirus* genus comprised of an AT-rich stretch and a portion of the GC-rich region. Altogether, this study provides a first insight into the mechanism by which anelloviruses manipulate host cell machinery to facilitate viral genome replication and represents a significant step forward in understanding the complex processes underlying anellovirus replication and persistent infection of these important commensal viruses.

## Introduction

Viruses of the family *Anelloviridae* are the dominant eukaryotic virus in the healthy human virome and are most abundant in the blood and bone marrow.^1–5^ Anelloviruses (ANVs) are non-enveloped, negative-sense, circular, single-stranded DNA (ssDNA) viruses that are nearly ubiquitous in human populations.^2,4^ Anelloviruses establish persistent infections from childhood, remaining detectable throughout life in greater than 90% of the global adult population.^6–9^ T cells are suspected to be the most likely reservoir of persistent ANV in humans;^10^ however, one study reported that granulocytes have the highest density of ANV genomes.^11^ The only known ANV pathogen is the chicken anemia virus (CAV), a gyrovirus which infects young chickens and induces anemia. CAV infects T cell precursors in the thymic cortex, CD8+ splenic lymphocytes, and hemocytoblasts in the bone marrow.^12^ Remarkably, with the exception of CAV, all other known ANVs are thought to be non-pathogenic and have evolved to coexist with their diverse vertebrate host species as commensal viruses.^6,13,14^ ANVs are not known to infect organisms outside of vertebrates.

Identification of Torque teno virus (TTV), the first ANV described, was reported in 1997.^15^ Since then, ANVs have been detected in a wide range of vertebrate species, including humans and other primates. Human ANVs are classified largely into three genera based on genome size: *Alpha-, Beta-,* and *Gammatorqueviruses*, corresponding to TTV (3.7-3.9 kb), Torque teno mini virus (TTMV) (2.8-2.9 kb), and Torque teno midi virus (TTMDV) (3.2 kb), respectively. More recently, another genus of human ANVs, *Hetorquevirus,* was recognized.^16,17^ Anelloviruses are the most prevalent eukaryotic virus found in humans and are highly genetically diverse, with thousands of unique capsid sequences identified across various tissue types.^2,6,13,17^ Additionally, it has been suggested that anelloviruses may benefit human health by shaping immunity during early development.^1,18–20^ A recent study reported that TTV infections induce an exhausted TTV-specific CD8+ T cell response which may mediate an imprinting of the immune system towards NKG2A+ T cells.^19^ It has been proposed that an immune system biased towards NKG2A^+^ CD8^+^ T cells is associated with protection against disease severity, mortality, and autoimmune/post-acute chronic disease.^19,21^ Importantly, elevated ANV levels in the blood are correlated with immunosuppression; thus, ANV levels in the blood have been proposed to serve as immune markers to inform clinical outcomes.^18,22^

Despite their near-universal presence in humans, there is limited understanding of the molecular virology of these ubiquitous yet enigmatic viruses. Viruses of the *Cressdnaviricota* phylum are circular Rep-encoding single-stranded DNA (CRESS DNA) viruses that replicate via a rolling-circle mechanism initiated by virus-coded Rep proteins of the HUH endonuclease superfamily.^23–25^ Owing to the absence of a recognizable Rep protein coding sequence, *Anelloviridae* is the only family of eukaryotic circular ssDNA viruses not included in this phylum.^23^ It has been proposed that the ANV origin of replication (ORI) is at a stem-loop structure with an octanucleotide motif that partially resembles the conserved nonanucleotide motif of CRESS DNA virus ORIs.^14,17,26^ However, there is no experimental evidence supporting this hypothesis. The protein coded by open reading frame 1 (ORF1) has been identified as the capsid protein (Cap) for ANVs, and it has also been suggested that the ORF1 protein may play a role in replication of the viral genome.^17,27–29^ Only recently has the ORF1 protein been experimentally demonstrated to form an ANV-like particle with icosahedral symmetry that is comprised of 60 jelly roll domain-containing protein subunits.^30^ Previous reports identified potential rolling circle replication (RCR) motifs associated with viral HUH Rep proteins as present in the ANV Cap.^29,31,32^ Recently, structural modeling of a TTMV Cap suggests the RCR motifs are spatially arranged in a manner that bears no resemblance to the HUH Rep proteins and that the highly conserved RNA-recognition-motif (RRM) fold does not form.^28^ In addition, these putative motifs are not well conserved in ANVs.^25^ The remaining ANV proteins lack homology to other known proteins, viral or otherwise;^14,33^ thus, their functions are largely unknown. The protein coded by ORF2 contains a conserved N-term W-x7-H-x3-C-x-C-x5-H motif which may suggest it has phosphatase activity; however, this activity has not been experimentally verified.^29,32,34,35^ Studying the function of ANV proteins has been challenging due to a lack of an *in vitro* cell culture system;^36^ although, recent advances in ANV virion production in MOLT-4 cells and the development of an ANV-based gene therapy vector system have paved the way for studying ANV protein functions.^37,38^

In this study, we report on the molecular mechanism of replication of human ANVs. Using a series of constructs to express viral gene products, we identified the protein coded by ORF2/3 as the human ANV replication initiation protein (Rip). Rip proteins from TTMV and TTV were necessary and sufficient to initiate replication from DNA constructs containing the entire non-coding regions (NCR) of either the *Beta-* or *Alphatorquevirus* genera, respectively. Further supporting this observation, we used immunoprecipitation-mass spectrometry (IP-MS) under multiple conditions to demonstrate that components of the polymerase alpha (POLα) and BTR complexes interact with ANV Rip during viral DNA replication, suggesting that ANVs may employ a recombination-dependent replication (RDR) mechanism for copying viral genomes. Additionally, we mapped the betatorquevirus ORI to a 92-bp sequence that immediately follows the protein coding sequences and contains an AT-rich stretch and a portion of the GC-rich region. To our knowledge, this is the first study to investigate and describe the molecular mechanism for human ANV replication and to experimentally demonstrate the function of the protein coded by ORF2/3.

## Results

### TTMV-X transcript mRNA3 accumulation is elevated at early times during wildtype virion production

The genomes of TTMVs are circular ssDNA that contain two distinct regions: the NCR and the protein coding sequences (Figure 1A). The NCR contains the viral promoter and all of the cis elements required for replication and packaging of viral genomes.^38^ The TTMV NCR comprises a GC-rich region with 2-5 highly conserved hairpin structures followed by conserved domain 1 (CD1). Downstream of CD1 is a TATA box and a hyper-conserved domain (HCD) which is conserved across the human ANV genera and serves as the transcriptional start site (Figure 1B). The proposed replication loop resides in conserved domain 2 (CD2).^17,26^ An intron is also contained within CD2 (Figure 1A and B). When this intron is spliced out, the Kozak sequence for ORF2 is formed. The TTMV coding region contains three known ORFs: ORF2, ORF1, and ORF3, in order from 5’ to 3’ (Figure 1A). Alternative splicing of the full transcript results in 8 detectable mRNA isoforms that code for up to 6 proteins (Figure 1B). The ORF2 and ORF1 proteins are coded by mRNA1; the ORF2/2 and ORF1/1 proteins are coded by mRNA2; the ORF2/3 and ORF1/2 proteins are coded by mRNA3 (Figure 1B).

**Figure 1.**
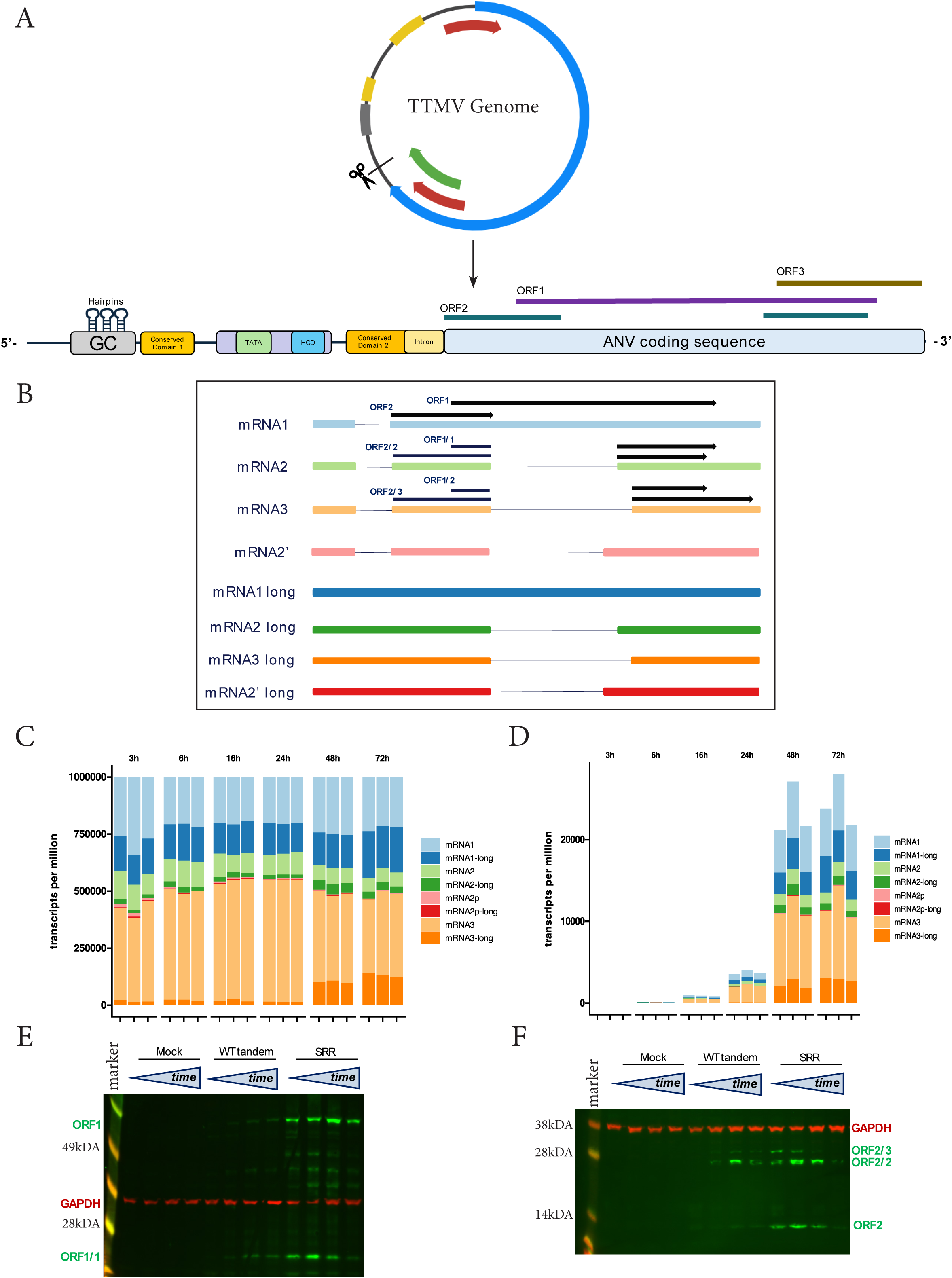
Relative viral mRNA transcript abundance and viral protein expression over time. (A) (Top) Circular representation of TTMV genome. (Bottom) A linearized representation of the TTMV genome annotated to represent key features of the coding and non-coding regions. The linearized sequence is represented as having the contiguous TTMV-NCR at the 5’ end of the genome and ending with the TTMV-X polyadenylation signal at the 3’ end. Sequence features of the NCR include the GC-rich region, which contains 3-5 highly conserved hairpin structures, conserved domain 1 (CD1), the promoter region (depicted in purple, containing the canonical TATA box and hyper conserved domain [HCD]), conserved domain 2 (CD2), and the UTR intron. Open reading frames are indicated with colored bars (ORF1 – purple; ORF2 – green; ORF3 – brown). Not to scale. (B) The eight detectable mRNA species of TTMV, which are generated via alternative splicing. Open reading frames are indicated by an ORF number and black arrow. (C-D) The relative abundance of the mRNA transcripts (B) was determined at specific timepoints during the viral life cycle using short-read RNA-seq. Relative expression of viral mRNAs was analyzed by normalization to (C) all viral reads or to (D) host + viral reads. Each bar represents one of three replicates per timepoint. (E-F) Immunoblots showing the detectable viral proteins expressed in MOLT-4 cells over 4 days using either WT Tandem or SRR constructs. Antibodies against (E) the C-term domain of ORF1 or (F) the N-term domain of ORF2 were used to detect viral proteins. GAPDH was used as a loading control.

DNA virus life cycles tend to follow a cascade of events, with early viral gene product functions biasing towards manipulation of host cell innate defenses and initiating viral genome replication. Viral gene products that are associated with packaging, virion assembly, and egress are usually expressed to higher levels following the onset of genomic replication.^39^ We used ribonucleic acid sequencing (RNA-seq) and immunoblotting with antibodies specific for either the C-terminal (C-Term) of ORF1 proteins or the N-terminal (N-term) of ORF2 proteins to investigate the kinetics of viral gene expression. Variants of mRNAs 1-3 in which the intron contained in CD2 is not spliced out were detected by RNA-seq (Figure 1B, C, and D). As the Kozak sequence for ORF2 does not form if this intron is not spliced out, it is not clear if these mRNA species are immature transcripts or if they are variants that bias towards translation from the ORF1 Kozak sequence.

We previously reported the development of an ANV particle production system that uses plasmids containing two tandem ANV genomes to express viral proteins.^37^ Interestingly, cells transfected with a plasmid containing tandem genomes of a TTMV identified from human tissue samples using the AnelloScope platform,^2,37^ here referred to as TTMV-X, accumulated increasing amounts of the “long” mRNA isoforms over time (Figure 1C and D). Another set of minor mRNA species, mRNA2’ and mRNA2’ long, were detected (Figure 1B-D). However, their functions are not yet known. Over a 72-hour time course of viral replication and virion production, mRNA3 appears to be the most abundant transcript, particularly at early time points (Figure 1C and D). The overall number of viral transcripts detected increased over time, peaking at 48 hours post-transfection (hpt) (Figure 1D). As with the RNA-seq, we used TTMV-X tandem genome constructs to monitor the expression of viral proteins over the course of 4 days (Figure 1E and F). Viral proteins translated from ORF1 and ORF1/1 accumulated over the course of 4 days with little to no detectable protein on day 1 to the peak accumulation on days 3-4 (Figure 1E). The levels of proteins translated from ORF2/3 and ORF2/2 peaked earlier, between days 2-3, while the protein coded by ORF2 peaked from days 3-4 (Figure 1F). We recently reported on the development of a self-replicating rescue plasmid (SRR) designed for expression of viral proteins to trans-rescue replication and packaging of ANV-based gene therapy vectors in MOLT-4 cells.^38^ The SRR expresses the simian virus 40 (SV40) large T antigen (LT) to drive replication of the plasmid through an SV40 ORI. This enables expression of a protein of interest from an upstream cassette in MOLT-4 cells, which are refractory to expression from exogenous DNA. Compared to the TTMV-X tandem genome construct, ANV protein expression from the SRR demonstrated accelerated expression kinetics and higher protein levels, with high levels of the protein coded by ORF2/3 occurring 1-day post-transfection followed by peak accumulation of ORF2 and ORF2/2 by 2 days post-transfection (Figure 1F). We currently do not have an antibody reagent capable of detecting the protein coded by ORF1/2. The early accumulation of mRNA3 and the proteins coded by ORF2/2 and ORF2/3 suggests they may play a role in early events of the ANV life cycle, such as innate immune modulation and viral DNA replication.

### The protein coded by TTMV-X ORF2/3 is necessary and sufficient to drive ANV reporter construct replication

We next investigated the potential for proteins coded by TTMV-X to drive replication of a reporter construct. Based on RNA-seq and protein data, we hypothesized that proteins translated from mRNA3 may play a role early in the ANV life cycle, including replication of the viral genome. To evaluate this hypothesis, we generated three constructs based on the SRR expression plasmid that contain the sequences of either the mRNA1, mRNA2, or mRNA3 transcripts. Each one of these transcripts can translate two proteins, one from the ORF2 frame and one from the ORF1 frame. We used these expression constructs to drive replication of a reporter construct that has the entire TTMV-X NCR driving an enhanced green fluorescent protein (eGFP) cassette. Using a Southern blot assay with a set of probes targeting the eGFP coding sequence, we detected DpnI-resistant DNA in cells transfected with the reporter plasmid and either the TTMV-X SRR or the SRR encoding mRNA3 (Figure 2A). DpnI targets the methylated adenine within GATC sequences of DNA that were replicated in *E. coli*, leading to digestion of the input DNA. In contrast, DNA replicated in eukaryotic cells remains resistant to DpnI.^40,41^ Cells transfected with SRRs encoding mRNA1 and mRNA2 did not produce DpnI-resistant DNA bands (Figure 2A). As mRNA3 can produce both ORF2/3 and ORF1/2 proteins, we used SRRs that encoded either protein to detect replicated DNA via a Southern blot assay. As previously observed, the mRNA3 expression construct produced a DpnI-resistant band of plasmid size (Figure 2B). Likewise, the ORF2/3 expression construct produced a DpnI-resistant band of plasmid size. In contrast, the expression construct for ORF1/2 did not produce a DpnI-resistant band. These observations suggest that the protein coded by ORF2/3 is the Rip protein for TTMV-X.

**Figure 2.**
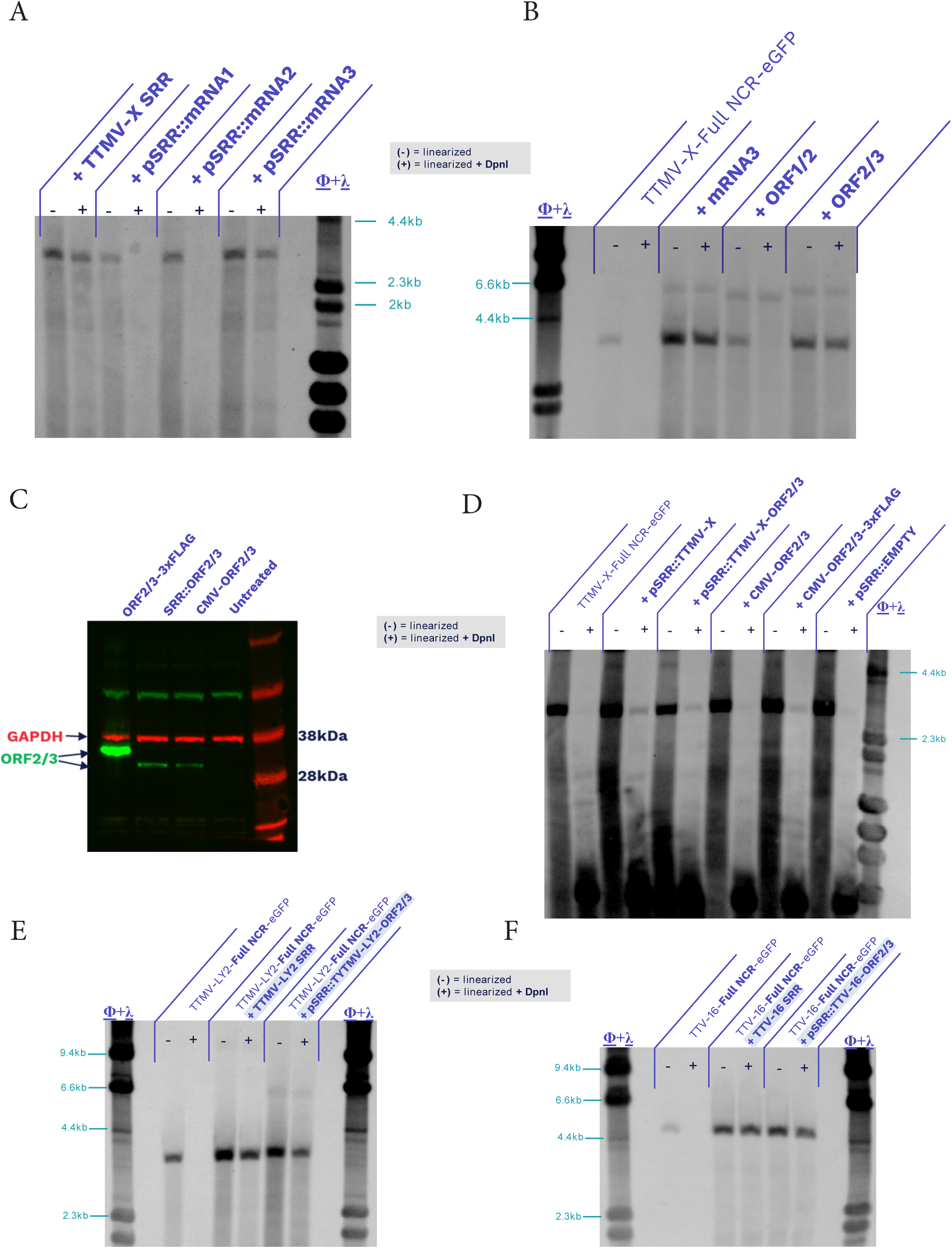
The ORF2/3 coded protein is necessary and sufficient for initiation of viral DNA replication. (A) Southern blot probing for eGFP DNA sequence in Day 3 lysates of MOLT-4 cells transfected with the TTMV -X-Full NCR-eGFP reporter and one of four separate SRR constructs expressing either the full TTMV-X coding sequence (TTMV-X SRR) or each of the three canonical TTMV mRNAs. Each condition was treated with a single-cut restriction enzyme to linearize the TTMV-X-Full NCR-eGFP sequence with (+) or without (-) DpnI. (B) Southern blot probing eGFP DNA sequences as in Fig.2C, but for MOLT-4 cells transfected with the TTMV-X-Full NCR-eGFP reporter alone or with constructs expressing mRNA3, ORF1/2, or ORF2/3. Each condition was treated with a single-cut restriction enzyme to linearize the TTMV-X-Full NCR-eGFP sequence with (+) or without (-) DpnI. (C) Immunoblot probing for the protein encoded by ORF2/3 when expressed by constructs expressing the ORF2/3 coding sequence in either an SRR format (+ SRR::ORF2/3) or in non-SRR formats with a CMV promoter driving expression of the ORF2/3 coding sequence (CMV-ORF2/3 and ORF2/2-3xFLAG). GAPDH was probed as a loading control. (D) Southern blot probing for eGFP sequences when TTMV-X-Full NCR-eGFP was transfected into HEK293 cells alone or with ORF2/3 expression constructs: pSRR::TTMV-X (full ANV coding sequence), pSRR::EMPTY (expresses no ANV viral proteins), CMV-ORF2/3, or CMV-ORF2/3-3xFLAG. (E) Southern blot probing for eGFP sequences when TTMV-LY2-Full NCR-eGFP was transfected into MOLT-4 cells alone or with expression construct coding the full TTMV-LY2 coding sequence (+ TTMV-LY2 SRR) or the protein encoded by ORF2/3 of TTMV-LY2 (+ pSRR::TTMV-LY2-ORF2/3). (F) Southern blot probing for eGFP sequences when TTV-16-Full NCR-eGFP was transfected into MOLT-4 cells alone or with expression construct coding the full TTV-16 coding sequence (+ TTV-16 SRR) or the protein encoded by ORF2/3 of TTV-16 (+ pSRR::TTV-16-ORF2/3).

Because the SV40 LT is a viral replication protein, we could not rule out that viral reporter replication observed using an SRR-based construct was driven by LT. While the SRRs for mRNA1, mRNA2, and ORF1/2 did not result in replicated reporter plasmid as detected by Southern blot, we set out to determine if the protein coded by ORF2/3 could replicate a reporter construct in the absence of LT. We generated two CMV-driven ORF2/3 expression plasmids that do not produce LT, with and without a 3x FLAG tag at the ORF2/3 C-term. Both the ORF2/3 and ORF2/3-3xFLAG constructs produce the ORF2/3 protein as detected by immunoblot (Figure 2C). While the ORF2/3 expression construct produced similar or slightly less protein than the SRR-ORF2/3 construct, the ORF2/3-3xFLAG construct produced a greater quantity of protein, possibly through stabilizing the protein (Figure 2C). Interestingly, the truncation of the last 69 amino acids of the C-term of ORF2/3 also resulted in an increased accumulation of protein (Supplemental Figure 1A). We used these expression constructs to transfect HEK293 cells that lack the LT protein. The ORF2/3 expression construct did not produce detectable DpnI-resistant DNA by Southern blot; however, the ORF2/3-3xFLAG construct did produce detectable DpnI-resistant reporter DNA in the absence of LT (Figure 2D). Taken together, these experiments demonstrate that the TTMV-X ORF2/3 protein acts as the viral Rip and is both necessary and sufficient to drive replication of DNA from the viral NCR.

### The protein coded by ORF2/3 is the Rip for TTMV-LY2 and TTV-16

Following identification of the TTMV-X ORF2/3 as the Rip responsible for driving replication from the viral NCR, we set out to test if the protein coded by ORF2/3 was also the Rip for other betatorqueviruses and for alphatorqueviruses. To test this, we first made reporter constructs that have an eGFP cassette driven by the NCR of a betatorquevirus identified from human pleural effusion, TTMV-LY2 (GenBank: JX134045.1)^42^ or a similar reporter construct with the entire NCR of TTV-16 (GenBank: AB017613.1),^43^ an alphatorquevirus (Supplemental Figure 1B). The NCR of alphatorqueviruses is approximately twice the size of the NCR of betatorqueviruses and contains two GC-rich stretches. We transfected the TTMV-LY2 reporter into MOLT-4 either alone, with a TTMV-LY2 SRR encoding all viral proteins, or a TTMV-LY2-ORF2/3 SRR that expresses only the ORF2/3 protein. DpnI-resistant reporter DNA was detected in both the TTMV-LY2 SRR and the TTMV-LY2-ORF2/3 SRR conditions (Figure 2E). We also observed that expression of eGFP from the LY2 NCR-driven reporter cassette was dependent on the presence of viral proteins (Supplemental Figure 1C). As observed with TTMV-LY2 constructs, the ORF2/3 of TTV-16 was capable of producing DpnI-resistant DNA from the construct containing the TTV-16 NCR (Figure 2F), and expression of the eGFP reporter cassette was dependent on the presence of viral proteins (Supplemental Figure 1D). Our results indicated that the proteins coded by ORF2/3 were both necessary and sufficient to drive DNA replication from the TTMV-LY2 and TTV-16 NCRs. These observations argue that the ORF2/3 protein serves as the Rip for human ANVs and possibly for ANVs infecting other vertebrates.

### The TTMV-X origin of replication maps to a 92bp sequence just downstream of the coding sequences

The ANV Rip was capable of activating the viral promoter to drive expression of an eGFP reporter, suggesting that either ORF2/3 can activate the promoter or that the promoter may activate through DNA replication (Supplemental Figure 1C and 1D). We aimed to determine how ANV proteins influence viral promoter expression. Constructs with deleted GC (ΔGC) or both GC and CD1 regions (ΔGC-ΔCD1) were designed to remove potential regulatory elements upstream of the TATA box (Supplemental Figure 2A). MOLT-4 cells were transfected with TTMV-X-Full NCR-eGFP, ΔGC, or ΔGC-ΔCD1, with or without TTMV-X mRNA3, ORF1/2, or ORF2/3 expression constructs. DNA was analyzed via Southern blot. As expected, TTMV-X-Full NCR-eGFP replicated in the presence of mRNA3 or ORF2/3, while ΔGC and ΔGC-ΔCD1 constructs showed no DpnI-resistant DNA, suggesting disrupted replication initiation (Supplemental Figure 2B and C). eGFP positivity was 50-70% for TTMV-X-Full NCR-eGFP and ΔGC but <10% for ΔGC-ΔCD1 (Supplemental Figure 2D). MFI analysis showed that both TTMV-X-Full NCR-eGFP and ΔGC constructs had elevated GFP levels with viral proteins, though ΔGC-ΔCD1 showed no increase (Supplemental Figure 2E). These results suggest that viral proteins enhance promoter expression without DNA replication, and the CD1 region is crucial for promoter activity. Interestingly, the results also suggest that GC region elements may be essential for replication from the ANV NCR.

While there is no experimental evidence to support it, it has been suggested that the ORI for ANVs lies within the CD2 region of the ANV NCR.^14,17,26^ The proposed octanucleotide motif bears some resemblance to the nonanucleotide CRESS DNA virus ORI motif and has been speculated to be part of a hairpin structure that may be cleaved by the previously undescribed replication protein(s), so it has also been referred to as the replication loop. However, the ANV Rip has no homology to the HUH Reps of CRESS DNA viruses. Additionally, we observed that a ΔGC was incapable of replicating DNA in the presence of ANV Rip (Supplemental Figure 2B), suggesting that elements in the GC region maybe play a role in replication. To evaluate the replication loop hypothesis, we designed a series of constructs based on the TTMV-X-Full NCR-eGFP reporter which have increasingly large truncations from the 3’ end of the NCR (Δ-1-5) (Figure 3A). Surprisingly, constructs Δ-1-4 produced DpnI-resistant bands as did the TTMV-X-Full NCR-eGFP control when transfected into MOLT-4 cells with the TTMV-X SRR (Figure 3B). The replication appeared to be less efficient when portions of the NCR were removed, consistent with results from truncated Porcine circovirus ORIs.^44^ Consistent with our observations from the ΔGC reporter, construct Δ-5, which also removes the GC-rich region, did not to produce DpnI-resistant DNA, indicating a failure to initiate replication. We next designed a series of constructs with truncations from the 5’ end of the contiguous TTMV-X NCR (Figure 3C). The smallest truncation, Δ-6, removes a 59nt sequence of DNA that lies between the ANV polyadenylation (polyA) signal and the GC-rich region. Unlike the TTMV-X-Full NCR-eGFP construct, all 5’ truncation constructs, Δ-6-9, failed to produce DpnI-resistant DNA when transfected with the TTMV-X SRR (Figure 3D). Together, these results argue that the ANV ORI resides in the 5’ end of the contiguous NCR rather than in the CD2 region.

**Figure 3.**
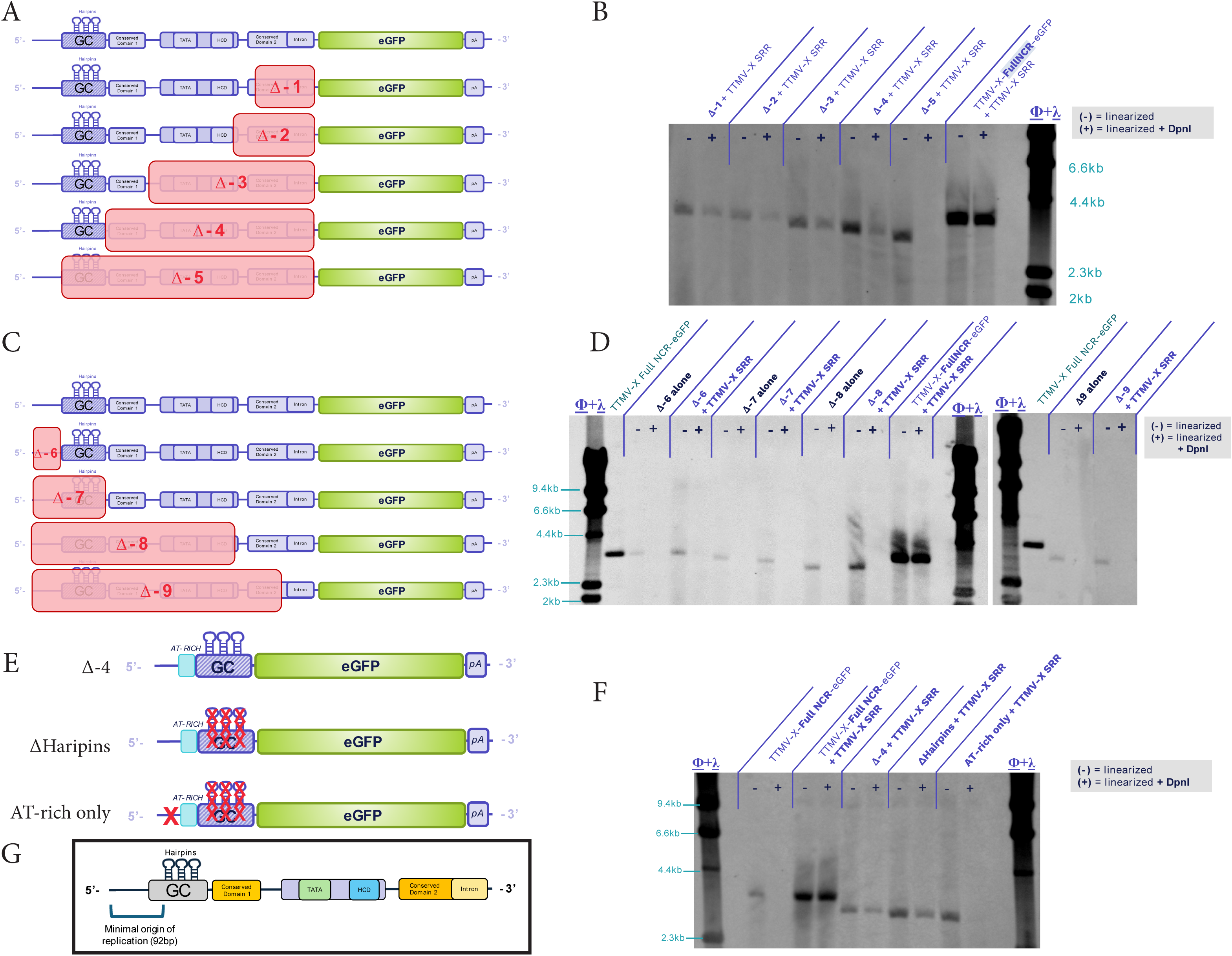
The TTMV-X minimal origin of replication maps to a 92nt sequence. (A) Linear representations of deletion mutants of TTMV-X-Full NCR-eGFP from the 3’ end of the NCR toward the 5’ targeting key sequence features (Δ-1-5). Targeted regions that were deleted are outlined in red and their respective nomenclature is denoted within each red box. (B) Southern blot probing for eGFP DNA sequences when the deletion mutants Δ-1-5 were co-transfected with an expression construct with the full ANV coding sequence (+ TTMV-X SRR). (C) Linear representations of deletion mutants of TTMV-X-Full NCR-eGFP from the 5’ end of the NCR toward the 3’ targeting key sequence features (Δ-6-9). Targeted regions that were deleted are outlined in red and their respective nomenclature is denoted within each red box. (D) Southern blot probing for eGFP DNA sequences when the deletion mutants Δ-6-9 were co-transfected with an expression construct with the full ANV coding sequence (+ TTMV-X SRR). (E) Linear representations of deletion mutants of the Δ-4 construct. The AT-rich near the GC-rich domain is depicted in blue. Deleted sequences are denoted by red Xs. (F) Southern blot probing for eGFP DNA sequences when Δ-4, ΔHairpins, or AT-rich only constructs were co-transfected with the full ANV coding sequence (+ TTMV-X SRR) in MOLT-4 cells. (G) A linear representation of the TTMV-X full NCR depicting the 92bp minimal origin of replication sequence.

The Δ-4 construct contained the minimal amount of TTMV-X NCR that was sufficient to replicate DNA with TTMV-X proteins (Figure 3B). This region contains several notable genetic elements which include a set of three highly conserved hairpin repeats in the GC-rich region and a 27nt AT-rich region (7% GC) immediately upstream of the GC-rich region, a common feature of ORIs for prokaryotes, eukaryotes, and DNA viruses.^45–49^ To map the TTMV-X minimal ORI, we made further modifications to the Δ-4 by deleting the conserved hairpins (ΔHairpins) and by deleting the hairpins and the sequence upstream of the AT-rich region (AT-rich only), leaving the AT-rich region along with the succeeding 34nt of GC-rich region upstream of the hairpins (Figure 3E). The TTMV-X-Full NCR-eGFP produced DpnI-resistant DNA only in the presence of the TTMV-X SRR, as expected (Figure 3F). Surprisingly, like the Δ-4 construct, the ΔHairpins construct produced DpnI-resistant DNA in the presence of the TTMV-X SRR, indicating that the highly conserved hairpins are not required for initiating replication from the ANV Ori. Loss of the sequence preceding the AT-rich region abolished DNA replication. The 92bp of TTMV-X sequence that comprises the ΔHairpins construct has very high homology to the corresponding sequence of TTMV-LY2 (Supplemental Figure 2F). These observations indicate the 92nt sequence immediately following the ANV polyA signal is sufficient to act as the ORI for betatorqueviruses (Figure 3G).

### TTMV-X Rip interacts with the DNA Polymerase α Complex and the BTR complex during genome replication

As the TTMV-X Rip has no homology to known proteins, including the Reps of CRESS DNA viruses, it is unknown how it might facilitate replication from the ANV ORI. To investigate how Rip might initiate replication, we performed IP-MS on MOLT-4 cells expressing a 3xFLAG-tagged TTMV-X Rip from an SRR. Rip-3xFLAG was immunoprecipitated with either a FLAG antibody (Ab) or an Ab targeting the ORF2 N-term domain. Gene ontology (GO) analysis revealed that proteins enriched in both the FLAG and ORF2 IP-MS were heavily involved in resolution of recombination intermediates and DNA synthesis (Figure 4A). Of particular interest, all four subunits of the DNA POLα complex^50^ (POLA1 catalytic subunit; POLA2 regulatory subunit; primase subunits PRIM1 and PRIM2) and the core components of the BTR complex^51–53^ (BLM RecQ helicase [BLM]; Topoisomerase 3A [TOP3A]; RMI1; RMI2) were detected in both pulldown schemes (Figure 4B-C). The POLα complex is the polymerase complex that is responsible for the priming of both the leading and lagging strands and initiation of DNA replication at ORIs.^54,55^ The BTR complex, also known as the BLM dissolvasome, regulates homologous recombination and facilitates dissolution of Holliday junctions.^53,56^ The BTR complex also plays a significant role in the alternative lengthening of telomeres (ALT) pathway and C-circle formation that are hallmarks of many cancers.^57–59^

The initial IP-MS experiment in MOLT-4 cells used constructs that expressed the LT to facilitate expression of proteins from plasmids. Additionally, lysates were not treated with nucleases so that we might capture indirect interactions mediated by nucleic acid. As a follow up experiment, we performed IP-MS in MOLT-4 cells and HEK293 cells, systems in which LT was not expressed, and in a HEK293T system that does express LT. All three experimental schemes incorporated DNase and RNase treatments to examine more direct interactions. Immunoprecipitation was performed using antibodies specific for the ORF3 domain of Rip. Components of the POLα complex (POLA1, POLA2, and PRIM1) and BTR complex (TOP3A and RMI1) were detected as interactors of TTMV-X Rip in all three cell lines, including HEK293 and MOLT-4 cells in which no LT was present (Figure 4D-G). In addition to POLα and BTR components, minichromosome maintenance (MCM) proteins 2-7 were detected in IP-MS from all three cell lines (Supplemental Figure 3A-C), but the enrichment was most significant in MOLT-4 and 293T cells lines. MCM proteins 2-7 are the six subunits of the MCM helicase complex which melts DNA strands at origins of replication.^60–62^ These IP-MS experiments support our observation that the TTMV-X ORF2/3 protein serves as the ANV Rip. Furthermore, the interaction with the BTR complex suggests that ANVs may use a recombination-related mechanism during replication.

### TTMV-X Rip protein modeling predicts Zn^2+^ coordination and potential dimerization with other viral proteins that share the ORF2 N-term domain

To gain more insight into the TTMV-X Rip, we used AlphaFold to model a putative protein structure.^63^ The N-term ORF2 domain of TTMV-X Rip is predicted, with very high confidence, to form 3 helices near the N-terminus; the first two helices possibly coordinating a Zn^2+^ ion (Figure 5A). The ORF3 portion of the protein is modeled to be largely unstructured, with a single helix near the C-term. The model also predicts possible interaction between N-term ORF2 domains of two TTMV-X Rip proteins, both coordinating Zn^2+^ (Figure 5B). While other ORF2 domain-containing proteins (ORF2 and ORF2/2) were not sufficient to initiate replication, they both contain the same N-term domain as the Rip protein. Thus, it is possible Rip interacts with other ORF2 domain-containing proteins (Figure 5C and D). We ran folding predictions on the TTMV-LY2 ORF2/3 protein and found the predicted structure to be very similar to that of the TTMV-X Rip (Figure 5E-G). This raises an intriguing possibility that interactions between the ORF2 domain-containing proteins could play a role in modulating replication processes.

**Figure 5.**
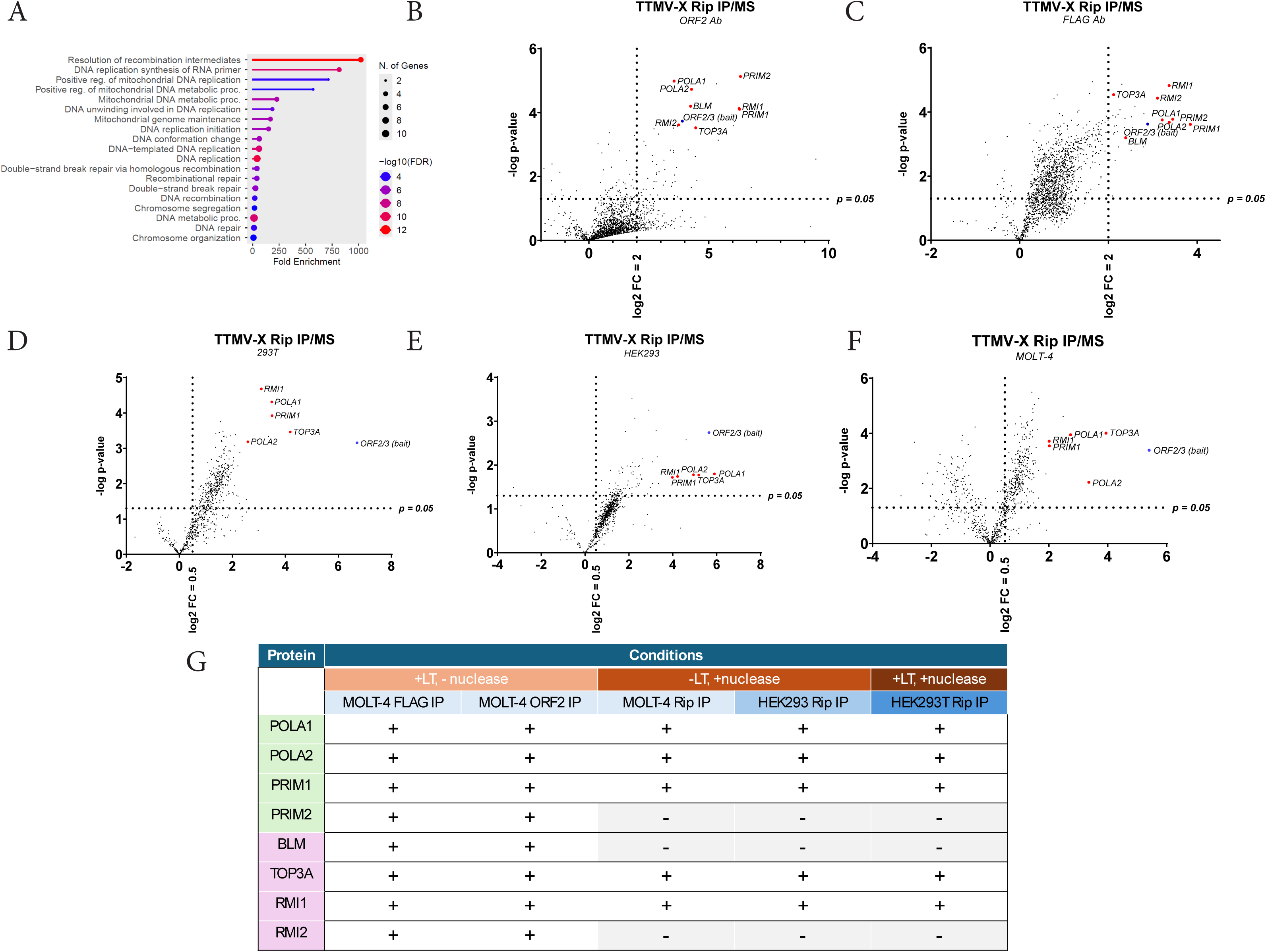
Rip IP/MS is enriched with proteins involved in DNA replication initiation and homologous recombination. (A) Overlapping hits between the FLAG and ORF2 IP/MS data sets with log2 FC > 2 & p < 0.05 were analyzed using the Go Ontology Biological Process algorithm. (B-C) ORF2 and FLAG IP/MS hits from Rip-expressing MOLT-4 cells are represented in volcano plots. Log2 FC represents the ratio of peptide counts from Rip+ lysates to peptide counts from Rip-lysates. The ORF2/3 bait protein is highlighted in blue, and proteins involved in the BTR and Polymerase α Complex are highlighted in red. (D-F) Volcano plots of Rip IP/MS hits from HEK293 (D), 293T (E), and MOLT-4 (F) cells. (G) Table showing BTR and Polymerase α Complex hits in 5 different IP designs. (+) denotes enrichment of log2 FC > 2 with statistical significance p < 0.05; (–) denotes lack of enrichment in the indicated IP design.

**Figure 6.**
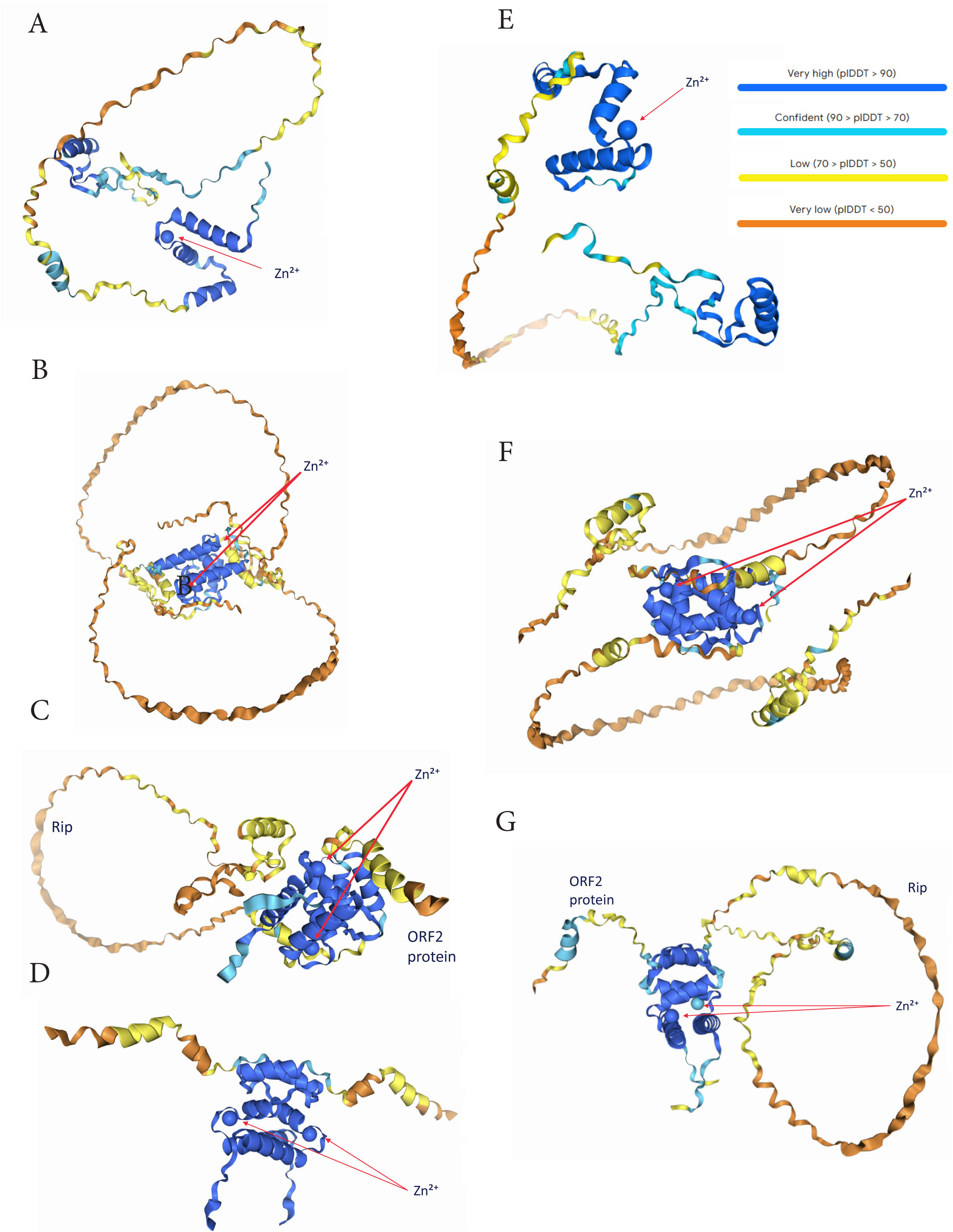
AlphaFold structural models predict TTMV-X Rip may act as a homo- or heterodimer with ORF2 domain containing proteins. (A) AlphaFold prediction of TTMV-X Rip folding in coordination with a Zn^2+^ ion (indicated with red arrows). TTMV-X Rip is predicted to (B) homodimerize with another Rip molecule, each coordinating Zn^2+^, or (C) for a heterodimer with another ORF2 domain containing proteins (here depicted as TTMV-X ORF2). (D) Prediction of TTMV-X ORF2 homodimer coordinating Zn^2+^ ions. (E) AlphaFold prediction of TTMV-LY2 Rip folding in coordination with a Zn^2+^ ion (indicated with red arrows). TTMV-LY2 Rip is predicted to (F) homodimerize with another Rip molecule, each coordinating Zn^2+^, or (G) for a heterodimer with another ORF2 domain containing proteins (here depicted as TTMV-LY2 ORF2). Prediction confidence is color coded – dark blue >90%, light blue 70-90%, yellow 50-70%, orange <50% confidence.

## Discussion

ANVs are exceptionally successful vertebrate viruses, and the near ubiquitous presence in humans as the dominant feature of the eukaryotic human virome makes it all the more surprising as to how little is known about their replication. In this study, we demonstrated that the ANV protein coded by ORF2/3 of both the *Betatorquevirus* and *Alphatorquevirus* genera, referred to as ANV Rip, is necessary and sufficient to initiate replication from cis elements located in the NCR region of the ANV genome. Furthermore, we identified components of both the POLα complex and the BTR complex as interacting with ANV Rip in multiple cell types, arguing that the ANV replication mechanism employs host cell machinery-mediated recombination.

While it has been assumed that ANVs would replicate using some form of RCR, no definitive evidence has been found for the presence of a Rep-like protein to facilitate initiation of RCR from the ANV genome. It had been speculated in the literature that the ORF1 protein, demonstrated to be the ANV capsid protein, contained RCR-related motifs that might resemble those found in Rep proteins.^29,31,32^ However, both sequence conservation analysis and predictive structural models called this hypothesis into question.^25,35^ Indeed, we found that the ORF1 protein is dispensable for driving replication from viral genetic elements. ANV Rip has no known homology to the HUH Rep proteins of CRESS DNA viruses or to any other known protein, suggesting ANVs may employ a novel mechanism for the replication of their circular ssDNA genomes.

We mapped the minimal TTMV-X ORI to a 92nt sequence of DNA that bears no resemblance to the replication loops of CRESS DNA viruses. The ANV ORI has an AT-rich stretch of DNA that is similar to the AT-rich stretches found at the ORIs of prokaryotes, eukaryotes, and dsDNA viruses.^46–49,55^ The SV40 ORI is embedded within its own promoter and has a 17bp AT-rich stretch.^46,66^ Herpes Simplex Virus (HSV) also has AT-rich stretches at its ORI.^47^ AT-rich sequences are often found at origins of DNA replication as the fewer hydrogen bonds between adenine and thymine facilitate the unwinding of the double-stranded helix of DNA.^46,48^ Unwinding of DNA at the ORI allows replication machinery to access the individual strands to prime and initiate replication. Interestingly, there is no evidence of a helicase domain or activity for ANV Rip. However, the detection of all six subunits of the MCM helicase complex argues that ANV Rip recruits the MCM helicase complex to melt open the ANV ORI. It is also unclear whether the ANV ORI initiates replication in a unidirectional or bidirectional manner. While RCR generally proceeds in a unidirectional manner, the ANV ORI may more closely resemble bidirectional cellular ORIs or the SV40 ORI than the replication loops of CRESS DNA viruses. It is also unknown whether the ANV ORI undergoes nicking, as observed for CRESS DNA virus RCR processes, but the presence of the MCM helicase complex proteins suggest ANV ORIs undergo a melting process to open the viral origin and initiate replication fork formation.

The detection of BTR complex components interacting with the ANV Rip offers some intriguing possibilities for modes of replication. The ALT pathway requires localization of the BTR complex to telomeres.^57^ The ALT pathway is a recombination-dependent form of telomere replication that is a hallmark of several forms of cancer, but it is also known to play a role in the replication and maintenance of some DNA virus genomes. ^67–70^ Many DNA viruses employ recombination strategies in the replication of their DNA,^71,72^ and ANVs are well documented to have recombination hotspots within their genomes.^2,17^ It is also of note that the BTR complex is required for the formation of extratelomeric ssDNA C-circles.^59,73^ It is possible the BTR complex plays a role in the formation of the genomic ssDNA in a manner that resembles the formation of C-circles. It is also interesting to note that TOP3A was detected in all IP-MS conditions. TOP3A has an isoform that localizes to the mitochondria and facilitates dissolution of hemi-catenated replication products after D-loop-mediated replication of the circular mitochondrial genome;^74^ thus, it is possible TOP3A may play a role in the dissolution of ANV replication intermediates. While additional experimentation is required to elucidate the role of TOP3A and the BTR complex in ANV replication, it is clear that host cell recombination machinery plays an important role in this process.

The AlphaFold models presented here suggest that the conserved ANV Rip may be largely unstructured throughout its ORF3 domain while the ORF2 domain of the Rip N-terminus is predicted to have helical structures that coordinate a Zn^2+^ ion. This zinc finger-like domain may serve to function as a mechanism for targeting the ANV origin, possibly while dimerized with another ORF2-domain containing protein. Interaction of different ORF2 domain-containing ANV proteins may even play a role in regulating phases of the ANV life. However, the functions of the other ORF2 domain-containing proteins remain to be explored. The ORF3 domain of Rip is located at the C-terminus and, other than a single helix, the AlphaFold model predicts this domain to be largely unstructured. Intrinsically disordered regions (IDRs) are often associated with compartments generated through liquid-liquid phase separation (LLPS);^64^ indeed, many viral proteins containing IDRs have been implicated in compartment formation through LLPS.^65^ This may aid the virus in establishing a replication compartment in which it can sequester proteins necessary for replication and packaging while potentially excluding restriction factors. We also observed that adding a 3xFLAG tag to or truncating the last 69 amino acids of the Rip C-term, resulted in enhanced accumulation of protein as detected by immunoblot. This suggests that the C-term of Rip may play a role in protein stability and turnover. Further studies will be needed to understand the properties of the domains within Rip and the roles they play in targeting the ANV ORI, recruiting host cell machinery, and protein stability.

## Methods and Materials

### Cell culture

MOLT-4 cells were sourced from the National Cancer Institute. These cells were cultured at 37°C with 5% CO2 in suspension culture using a complete growth medium (Gibco’s RPMI 1640 with 10% fetal bovine serum, supplemented with 1 mM sodium pyruvate, 0.1% Pluronic F-68, and 2 mM L-glutamine) while shaking at 100 rpm with over 85% relative humidity (RH). HEK293 and HEK293T cells were obtained from the American Type Culture Collection (Manassas, VA) and maintained in Dulbecco’s Modified Eagle’s Medium (DMEM; Corning, Corning NY) supplemented with 10% (v/v) fetal bovine serum (FBS) in humidified incubators at 5% CO2 and 37°C. Cells were regularly passaged by treatment with trypsin-EDTA (0.05%; Corning) and transferred to fresh media.

### Transfections

MOLT-4 cells were subjected to electroporation in 2S buffer (comprising 5 mM KCl, 15 mM MgCl2, 15 mM HEPES, 150 mM Na2HPO4 at pH 7.2, and 50 mM sodium succinate) using a NEPA electroporator. Cell pellets, containing approximately 1 × 10^7 cells, were resuspended in 500 µL of the 2S buffer along with 50 µg of expression plasmid and 50 µg of eGFP reporter plasmid, in amounts sufficient for five cuvettes with identical DNA content. Following electroporation, approximately 300 µL of pre-warmed media was added to each cuvette using a bulb pipette, and the cells were gently mixed to dissociate any clumps. The cell suspension was subsequently transferred to 25 mL of pre-warmed media in a flask. Flasks were incubated for three days at 37°C with shaking at 125 rpm. Prior to harvesting, cells were imaged using an EVOS M5000 microscope (Thermo Fisher, AMF5000SV) in both brightfield and GFP channels at 10x magnification. HEK293 and HEK293T cells were transfected using PEIpro (Polyplus) per manufacturer’s protocol. Cells were incubated for the indicated time at 37°C.

### RNA-Seq

Total RNA was extracted from transfected cells using Trizol (Thermo Fisher Cat# 15596026) per manufacturer’s protocol followed by rRNA depletion. NEBNext Ultra II Directional RNA Library Prep For Illumina (NEB Cat# E7760L) was used to prepare the sequencing library from the rRNA-depleted RNA samples per manufacturer’s protocol. Samples were sequenced with an Illumina NextSeq. Sequencing reads were processed using nf-core/rnaseq v3.17.0 (doi: 10.5281/zenodo.1400710) of the nf-core collection of workflows,^75^ utilizing reproducible software environments from the Bioconda^76^ and Biocontainers^77^ projects. We used the “star_salmon” workflow and chose fastp v0.23.4 for read quality filtering, while otherwise retaining the default settings and software versions used in nf-core/rnaseq v3.17.0. For the genome reference, we used the ENSEMBL v113 Homo sapiens genome, together with the genome sequence of TTMV-X, plus their respective annotations. We used the resulting alignments form the RNA-seq pipeline to quantify the host and TTMV-X transcript isoforms using StringTie v2.2.3,^78^ in units of Transcripts per Million Mapped (TPM), then extracted the TTMV-X transcripts using samtools v1.20^79^ to quantify the relative TPM values of the TTMV-X transcripts exclusively.

### Immunoblotting

Cells were harvested 3 days post-electroporation, washed with PBS, and lysed in 0.5% Triton-X, 300 mM NaCl, and 50 mM Tris pH 8.0. Genomic DNA was degraded using mSAN, and proteins were denatured with LI-COR protein sample loading buffer and reducing agent at 95°C for 10 minutes. Lysates were run on BOLT SDS-PAGE gels (12% for ORF2 and 4-12% for ORF1). Proteins were transferred to nitrocellulose membranes using the Bio-Rad Trans-Blot Turbo Transfer system. Membranes were blocked for 1 hour with LI-COR Intercept Blocking buffer, followed by overnight primary antibody incubation (1:1000) at 4°C. GAPDH detection used an antibody from Cell Signaling (Cat#97166S), while ORF1 and ORF2 used GenScript antibodies. Membranes were then incubated with IRDye® 800CW and 680CW goat IgG secondary antibodies (1:10,000) from LI-COR for 2 hours and imaged using the Bio-Rad ChemiDoc MP imaging system.

### Southern assay

For Southern Blot analysis, total nucleic acid from transfected MOLT-4 cells was extracted using the DNeasy Blood and Tissue Kit (QIAgen #69504) with proteinase K treatment. The DNA was linearized by overnight digestion at 37°C with a restriction endonuclease specific to the viral DNA. To digest input plasmid DNA, samples were treated with DpnI (NEB #R0176), which cleaves methylated GATC sites. Following digestion, 10 µg of DNA per lane was electrophoresed on a 1.0% agarose gel, then depurinated, denatured, and transferred overnight onto a Hybond-N+ membrane (Cytiva #RPN203B). The membrane was pre-hybridized for 4 hours in ULTRAhyb buffer (Thermo Fisher #AM8670) and probed overnight with biotin-labeled oligos specific to viral DNA. Probes were generated using the BioPrime™ Array CGH Genomic Labeling System (Thermo Fisher #18095011). Membranes were blocked for 30 minutes, incubated with IRDye800CW Streptavidin (LI-COR #926-32230) for 30 minutes, and imaged using the Bio-Rad ChemiDoc MP Imaging System.

### Flow Cytometry

The entire culture of cells was harvested by pipetting into 50mL conical tube at 3 days post transfection. 300uL of cells were transferred to 96-well round bottom plates. A Cytek Guava clow cytometer was used to measure %GFP and mean fluorescence intensity over 10,000 events.

### IP/MS

For immunoprecipitation, HEK293, HEK293T and MOLT-4 cells were co-transfected in triplicate with constructs expressing TTMV-X ORF2/3 or ORF2/3-FLAG and constructs containing the TTMV-X NCR region to generate Bait (+) lysates. Bait (-) lysates were generated by transfecting cells with the NCR construct alone. Cells were harvested 2 days post-transfection and washed once with PBS. Lysis was performed at 4C for 30 minutes in non-denaturing buffer (0.5% Triton-X, 300 mM NaCl, 50 mM Tris pH 8.0, protease/phosphatase inhibitor) with or without mSAN and RNase A. mSAN/RNase A-treated lysates were additionally incubated for 1 hour at 37C. Cell debris was pelleted at 4C for 15m at 10,000g and the clarified supernatants were subjected to IP using bead-conjugated antibodies targeting the C-term FLAG tag, the ORF2 Zn finger domain or the ORF2/3 C-term domain. ORF2 and ORF2/3 antibodies were conjugated to M-270 Epoxy beads using the Dynabeads™ Antibody Coupling Kit (Thermo Fisher #14311D) and FLAG IPs were performed using ANTI-FLAG® M2 Magnetic Beads (Sigma # M8823). Cell lysates were incubated with antibody-bead slurries overnight at 4C rotating end-over-end. Beads were pelleted using a MagStrip and washed twice with cold lysis buffer and proteins were eluted from the beads using 4X Protein Sample Loading buffer (LI-COR). Beads were incubated in elution buffer at 65C for 20 minutes and subsequently separated from the eluate using a MagStrip and stored at -80 °C until quantitative proteomic sample preparation and analysis (performed at IQ Proteomics, Cambridge, MA). Fold change values represent the ratio of Bait+ signal to Bait-signal and p-values were calculated using two-sample t-test.

### AlphaFold Server

Protein sequence from TTMV-X and TTMV-LY2 were input into the AlphaFold server (https://alphafoldserver.com/). pLDDT is a per-atom confidence estimate on a 0-100 scale where a higher value indicates higher confidence. *This information is subject to AlphaFold Server Output Terms of Use found at <u>alphafoldserver.com/output-terms</u>*.

## Acknowledgements

We would like to thank Brian Luque, Chris McNulty, and Kelly Morgan for facilitating this manuscript.

## Author Contributions

Conceptualization, JC, NB, ST, and DV; Methodology, JC, NB, ST, CP, and CE; Formal Analysis, JC, NB, ST, CE, NS, and PM; Experimentation, NB, ST, CD, JM, NS, and KT; Writing – Original Draft, JC, NB, and ST; Writing – Review & Editing, JC, NB, and ST; Visualization, JC, NB, ST, CE, and PJ; Supervision, JC and GP

## Supplemental Figure Legends

**Supplemental Figure 1. ORF2/3 is the replication protein of both *Beta-* and *Alphatorqueviruses*.** (A) Immunoblot probing detectable viral proteins expressed in MOLT-4 cells 3 days post-transfection using either TTMV-X SRR or TTMV-X SRR with an ORF2/3 truncation construct. Antibody against the N-term domain of ORF2 was used to detect viral proteins. Anti-GAPDH antibody was used as a loading control. (B) Linear representations of (Top) TTMV-LY2 reporter cassette containing the full NCR of TTMV-X followed by an eGFP coding sequence and SV40 polyA (TTMV-LY2-Full NCR-eGFP) and of (bottom)TTV-16 reporter cassette containing the full NCR of TTV-16 followed by an eGFP coding sequence and SV40 polyA (TTV-16-Full NCR-eGFP) (C) Fluorescence microscopy of conditions in (Figure 2E) viewed at a 10X magnification. (D) Fluorescence microscopy of conditions in (Figure 2F) viewed at a 10X magnification.

**Supplemental Figure2. CD1 of the TTMV-X is important for activation of the viral promoter.** (A) Linear representation of deletion mutants of TTMV-X-Full NCR-eGFP. Regions upstream of the core promoter that were deleted are outlined in the red boxes along with key feature deletions denoted inside (ΔGC and ΔGC-ΔCD1). Construct nomenclature is respective to their deletions. (B-C) Southern blots probing for eGFP sequences when the constructs depicted in (A) were transfected into MOLT-4 cells either alone or with SRR constructs expressing the proteins encoded by TTMV-X ORF1/2 (+ ORF1/2), ORF2/3 (+ ORF2/3), or both (+ mRNA3). (D-E) Flow cytometry data for eGFP expression for transfection conditions in (B-C). (D) The percentage GFP positive data (%GFP+) coupled with each condition’s respective (E) mean fluorescence intensity (MFI) data (log scale). (F) The 92nt ORI of TTMV-X was aligned against the same region of TTMV-LY2. Areas of sequence homology are highlighted in yellow. C-to-G or G-to-C variations are highlighted in blue. A-to-T or T-to-A variations are highlighted in green.

**Supplemental Figure 4. Rip IP/MS is enriched with MCM proteins.** (A-C) Volcano plots of Rip IP/MS hits from MOLT-4 (A), HEK293 (B), and 293T (C) cells with the MCM family proteins highlighted in red and the ORF2/3 bait highlighted in blue. The left panel shows every data point from the IP/MS experiment, and the right panel shows only the MCM data points for clarity.

