## Supplementary figures and images for "Anellovirus protein coded by ORF2/3 recruits host cell replication and homologous recombination machinery during replication"

### Supplemental figures

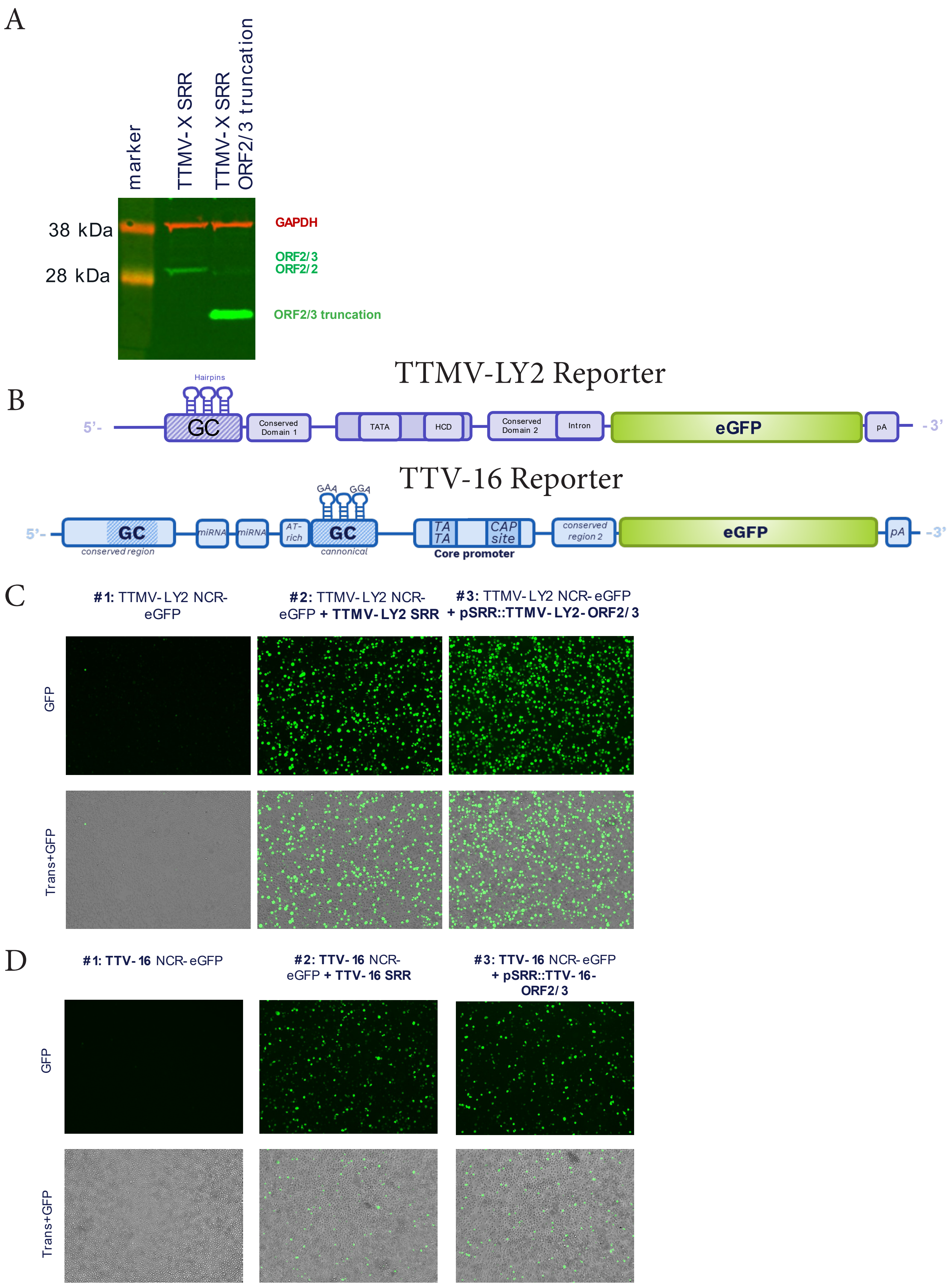

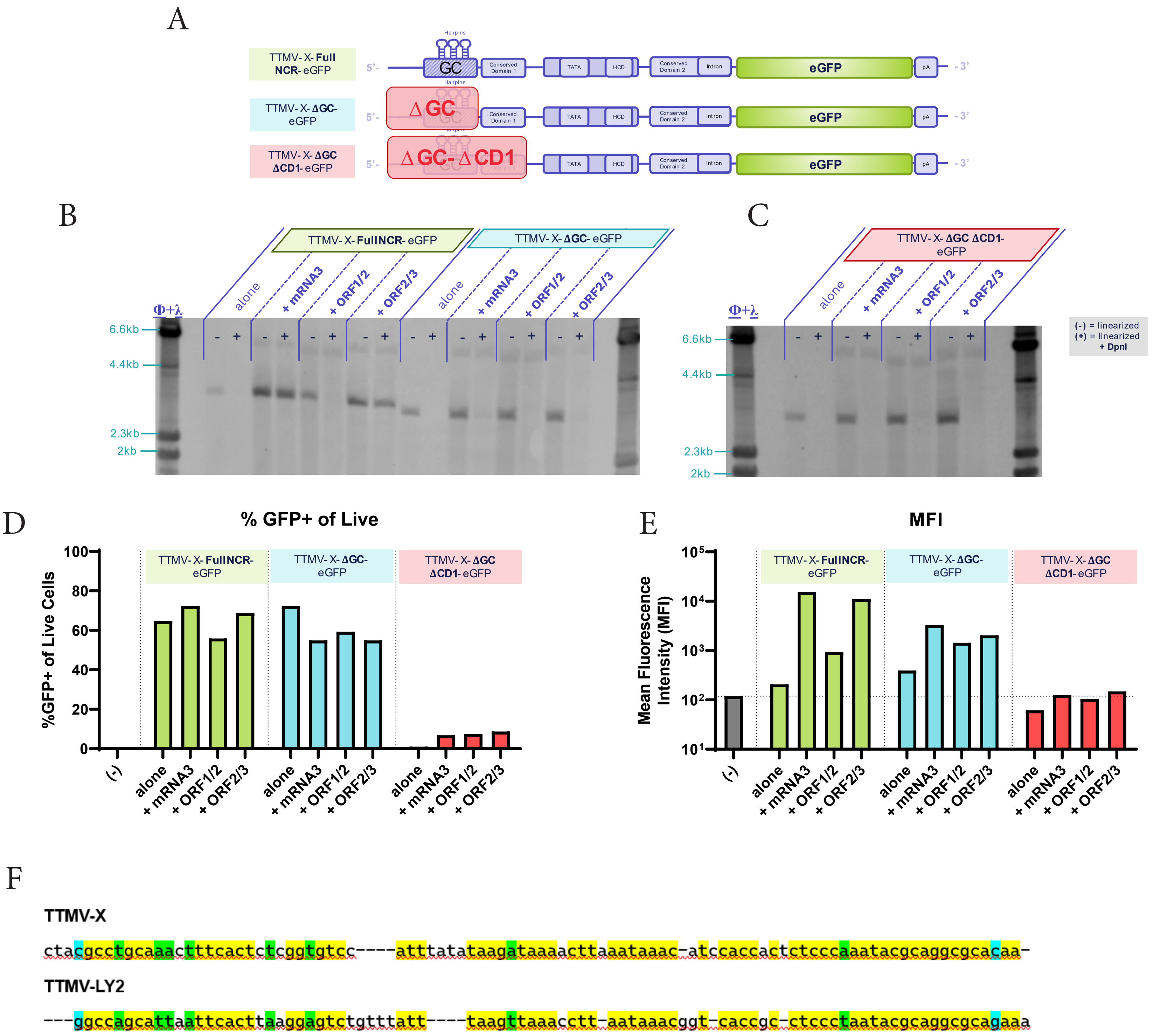

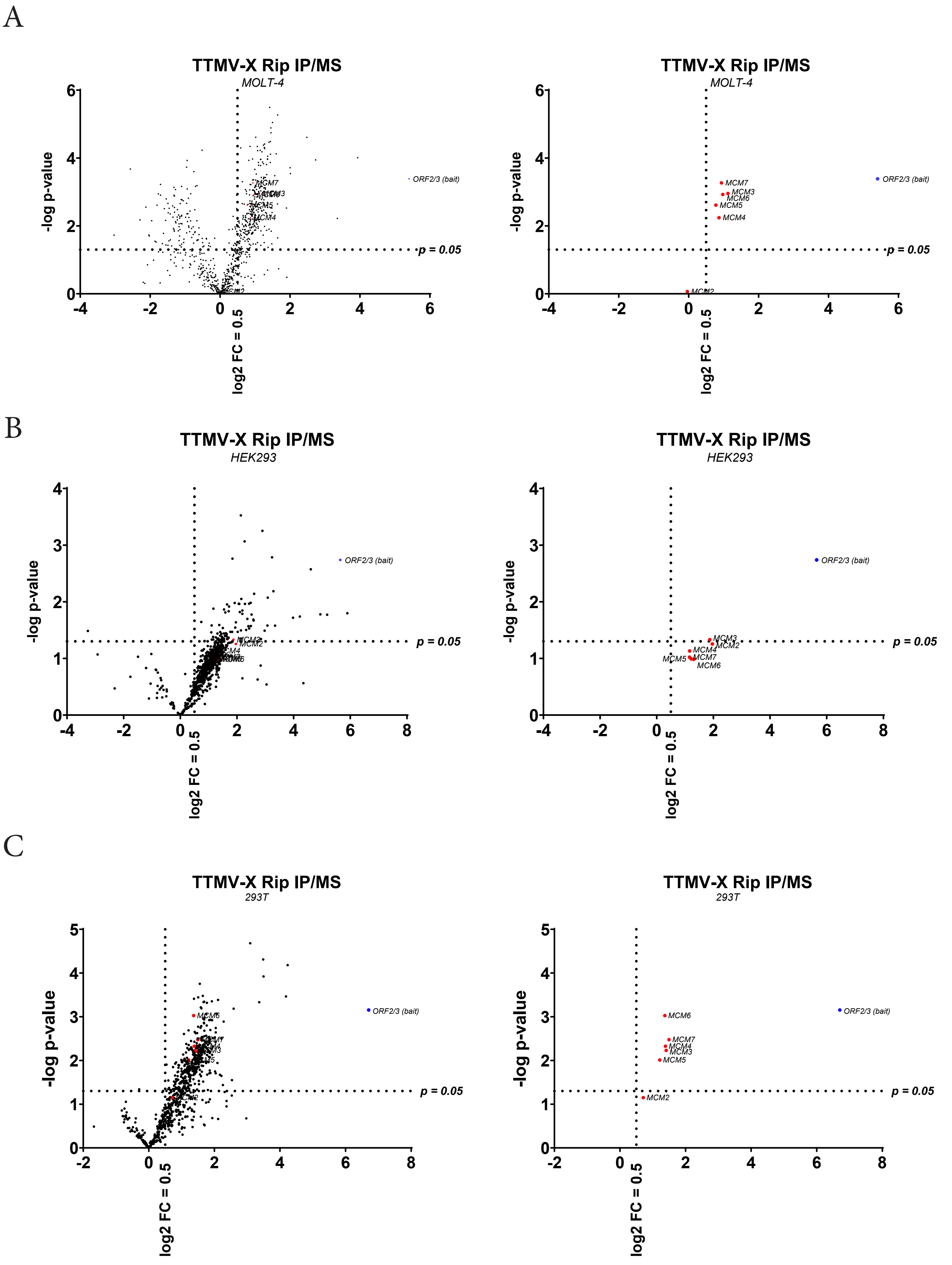
